# Development of a novel hybrid alphavirus-SARS-CoV-2 particle for rapid *in vitro* screening and quantification of neutralization antibodies, viral variants, and antiviral drugs

**DOI:** 10.1101/2020.12.22.423965

**Authors:** Brian Hetrick, Sijia He, Linda D. Chilin, Deemah Dabbagh, Farhang Alem, Aarthi Narayanan, Alessandra Luchini, Tuanjie Li, Xuefeng Liu, Joshua Copeland, Angela Pak, Tshaka Cunningham, Lance Liotta, Emanuel F. Petricoin, Ali Andalibi, Yuntao Wu

**Affiliations:** National Center for Biodefense and Infectious Diseases, School of Systems Biology, George Mason University, Manassas, VA 20110, USA; Center for Applied Proteomics and Molecular Medicine, George Mason University, Manassas, VA, 20110, USA; Department of Pathology, Center for Cell Reprogramming, Georgetown University Medical Center, Washington DC, 20057, USA; Department of Oncology, Lombardi Comprehensive Cancer Center, Georgetown University Medical Center, Washington DC, 20057, USA; TruGenomix Inc. 155 Gibbs Street, Room 559, Rockville, MD 20850, USA

## Abstract

Timely development of vaccines and antiviral drugs is critical to control the COVID-19 pandemic ^1–6^. Current methods for quantifying vaccine-induced neutralizing antibodies involve the use of pseudoviruses, such as the SARS-CoV-2 spike protein (S) pseudotyped lentivirus^7–14^. However, these pseudoviruses contain structural proteins foreign to SARS-CoV-2, and require days to infect and express reporter genes^15^. Here we describe the development of a new hybrid alphavirus-SARS-CoV-2 (Ha-CoV-2) particle for rapid and accurate quantification of neutralization antibodies and viral variants. Ha-CoV-2 is a non-replicating SARS-CoV-2 virus-like particle, composed of SARS-CoV-2 structural proteins (S, M, N, and E) and a RNA genome derived from a fast expressing alphavirus vector ^16^. We demonstrated that Ha-CoV-2 can rapidly and robustly express reporter genes in target cells within 3-6 hours. We further validated Ha-CoV-2 for rapid quantification of neutralization antibodies, viral variants, and antiviral drugs. In addition, as a proof-of-concept, we assembled and compared the relative infectivity of a panel of 10 Ha-CoV-2 variant isolates (D614G, P.1, B.1.1.207, B.1.351, B.1.1.7, B.1.429, B.1.258, B.1.494, B.1.2, B.1.1298), and demonstrated that these variants in general are 2-10 fold more infectious. Furthermore, we quantified the anti-serum from an infected and vaccinated individual; the one dose vaccination with Moderna mRNA-1273 has greatly increased the anti-serum titer for approximately 6 fold. The post-vaccination serum has also demonstrated various neutralizing activities against all 9 variants tested. These results demonstrated that Ha-CoV-2 can be used as a robust platform for rapid quantification of neutralizing antibodies against SARS-CoV-2 and its variants.

## INTRODUCTION

Severe acute respiratory syndrome coronavirus 2 (SARS-CoV-2) is a rapidly spreading, novel beta-coronavirus that is causing the ongoing global pandemic of coronavirus disease 2019 (COVID-19) ^17–21^. SARS-CoV-2 has caused over 140 million infections and 3 million deaths globally as of April 2021. Antiviral drugs and neutralizing antibodies are effective to combat the pandemic. In particular, neutralizing antibodies, induced by vaccines or by the virus, can play a critical role in controlling and preventing infection. Currently, only one FDA-approved drug, remdesivir, is available to reduce hospital stay ^1^; several vaccines have recently shown significant results in phase III clinical trials ^2–6^, and been approved for emergency use. Nevertheless, the effectiveness of vaccines needs to be continuously monitored for the induction of neutralizing antibodies against evolving viral variants.

Current antiviral drug screening and quantification of neutralizing antibodies rely on the use of SARS-CoV-2 pseudoviruses ^7–14^. The use of live virus requires biosafety level (BSL) 3 facility and practice, which limits large-scale testing and analyses in common laboratories. Both lentivirus and vesicular stomatitis virus (VSV), pseudotyped with the SARS-CoV-2 S protein, are used in cell-based neutralization assays and in antiviral drug screening ^8–11^. SARS-CoV-2 contains four structural proteins: the spike protein (S), the membrane protein (M), the envelope protein (E), and the nucleocapsid protein (N) ^22,23^. S is the major viral protein responsible for virus attachment and entry to target cells ^24–26^, and thus, is commonly used to pseudotype viruses. Nevertheless, both VSV- and lentiviral-based pseudoviral particles contain only the S protein, and the major viral structural components are foreign to SARS-CoV-2, which may affect viron properties in receptor binding and responses to antibody neutralization ^15^. In addition, an important issue for the VSV-based pseudovirus is the presence of residual VSV virus, which can result in high rates of false-positive results ^27^. Furthermore, the use of lenti-pseudoviruses for neutralization assay is time consuming, and requires 2 to 3 days to infect and generate reporter signals ^7–11^.

To overcome the limitations of current pseudoviruses, here we describe the development and validation of a new hybrid alphavirus-SARS-CoV-2 particle (Ha-CoV-2) for rapid quantification of neutralization antibodies and antiviral drugs. Ha-CoV-2 is a non-replicating SARS-CoV-2 virus-like particle that is composed of authentic virus structural proteins (S, M, N, and E) from SARS-CoV-2 with no structural proteins from other virus. Ha-CoV-2 also contains a genome derived from an alphavirus-based vector ^16, 28^, which can rapidly and robustly express reporter genes within a few hours after viral entry ^28^. In this study, we further demonstrate that Ha-CoV-2 can be used as a robust platform for rapid quantification of neutralization antibodies, viral variants, and antiviral drugs.

## RESULTS

To establish a rapid cell-based SARS-CoV-2 infection system for screening and evaluation of neutralizing antibodies and antiviral drugs, we developed a new hybrid alphavirus-SARS-CoV-2 viral particle, in which an alphavirus-based RNA genome is enclosed for rapid expression of reporter genes in target cells (**Fig. 1A**). The genomic RNA consists of the 5’ untranslated region and open-reading frames coding for the nonstructural proteins (nsp) 1-4 from Semliki Forest virus (SFV) ^16,29^; the inclusion of nsp1-4 allows for self-amplification of the RNA genome in cells. The RNA genome also contains viral subgenomic RNA promoters for the expression of reporter genes (such as luciferase). The 3’ end of the genome contains the 3’ untranslated region of SFV and a poly(A) tail that are used to stabilize RNA. In addition, a putative SARS-CoV-2 packaging signal was inserted downstream of the reporter gene to facilitate RNA packaging by the SARS-CoV-2 structural protein N. To assemble viral particles, we used the DNA vector, Ha-CoV-2, to express the genomic RNA. The Ha-CoV-2 vector was cotransfected with vectors expressing the structural proteins of SARS-CoV-2 (S, M, E, and N) into HEK293T cells (**Fig. 1A**). Virion particles were harvested at 48 hours post cotransfection, and tested for virion infectivity and the ability to express reporter genes in target cells. First, to confirm the presence of the SARS-CoV-2 structural proteins in Ha-CoV-2 particles, we performed western blots of purified particles, using antibodies against the S protein of SARS-CoV-2. We were able to detect the presence of S in Ha-CoV-2 particles (**Fig. 1B**). To further determine the presence of the other structural proteins of SARS-CoV-2, we also assembled Ha-CoV-2 particles using FLAG-tagged M and N proteins, and performed western blots using anti-FLAG antibodies. We were able to confirm the presence of both M and N in Ha-CoV-2 particles (**Fig. 1C** and **1D**). Furthermore, to determine whether these structural proteins are present in the same virion particles, we used anti-S antibody-conjugated magnetic beads to pull down Ha-CoV-2 particles. The magnetically separated particles were further probed with western blot using the anti-FLAG antibody for the presence of FLAG-tagged M protein. As shown in **Fig. 1E**, we detected the presence of M in the anti-S antibody pull-down particles, confirming that cotransfection of the SARS-CoV-2 structural proteins with the Ha-CoV-2 vector led to the production of SARS-CoV-2 virus-like particles (VLPs).

**Fig. 1.**
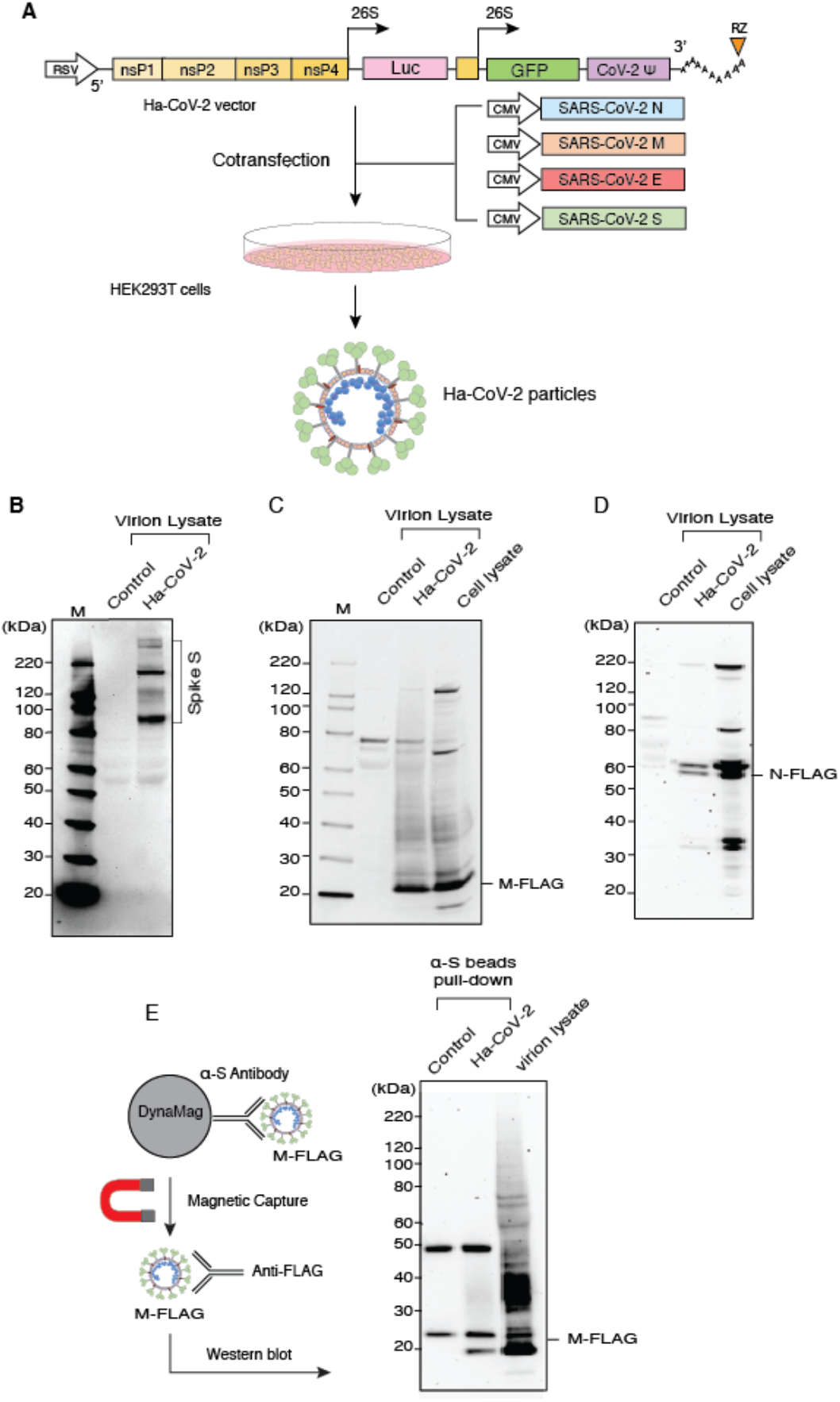
Design and assembly of Ha-Cov-2 particles. **(A)** Illustration of the design of Ha-CoV-2 vector. The vector contains a RSV promoter that transcribes the full-length viral RNA genome to be packaged into Ha-CoV-2 particles. Shown are the 5#8217; untranslated region followed by open-reading frames coding for nonstructural proteins (nsp) 1-4 from Semliki Forest virus (SFV), viral subgenomic promoters for Luc and GFP reporter expression, the 3#8217; untranslated region and a poly(A) tail that is self-cleaved by the hepatitis delta virus ribozyme (RZ). The SARS-CoV-2 packaging signal is inserted in front of the 3#8217; untranslated region. To assemble viral particles, HEK293T cells were co-transfected with Ha-CoV-2 and the vectors expressing the 4 structural proteins of SARS-CoV-2 (S, M, E, and N). HA-CoV2 particles in the supernatant were harvested at 48 hours, purified, lysed, and then analyzed by western blot using antibodies for the SARS-CoV-2 S protein (**B**). Control is the supernatant from cells transfected with the Ha-CoV-2 vector alone. (**C** and **D**) Particles were also assembled using FALG-tagged M and N. Particles were analyzed with western blot using an antibody against FLAG. **(E)** Particles in the supernatant were also captured with magnetic beads conjugated with the anti-S antibody, and then analyzed with western blot using the antibody again FLAG for FLAG-tagged M protein in the particles.

To further demonstrate the ability of Ha-CoV-2 particles to infect and express reporter genes in target cells, we assembled an Ha-CoV-2(GFP) reporter virus, and used it to infect HEK293T(ACE/TEMPRSS2) cells that overexpressed both ACE2 and TMPRSS2. We observed GFP expression in cells following infection (**Fig. 2A**), demonstrating that the alphavirus-based RNA genome can be packaged by the budding SARS-CoV-2 VLPs, and is capable of expressing the GFP reporter gene in target cells. To determine whether the infection of target cells by Ha-CoV-2 is dependent on the interaction of S with the ACE2 receptor ^18,30^, we assembled an Ha-CoV-2(Luc) reporter virus and used it to simultaneously infect HEK293T(ACE/TEMPRSS2) cells and the parent HEK293T cell. As shown in **Fig. 2B**, Ha-CoV-2(Luc) expressed high-levels of Luc in the HEK293T(ACE/TEMPRSS2) cells but minimal levels of Luc in HEK293T, demonstrating the requirement of ACE2 for Ha-CoV-2 infection. We further confirmed the requirement for S-ACE2 interaction by generating particles without the S protein. As shown in **Fig. 2C**, in the absence of S, Luc expression was highly diminished, demonstrating the requirement of S for Ha-CoV-2 infection. We also tested the requirement for the other structural proteins, M, N, and E, for Ha-CoV-2 infection. Although these proteins were found to be nonessential, removal of M, N, and E led to a reduction in Ha-CoV-2 infection (**Fig. 2C**). We further investigated the individual contributions of M, N, and E for Ha-CoV-2 infection. It appeared that in general, particles assembled with two or three of these structural proteins gave rise to a higher level of infection than those with only one protein. However, the presence of S plus E appears to be sufficient for the full infectivity of Ha-CoV-2 (**Fig. 2D**).

**Fig. 2.**
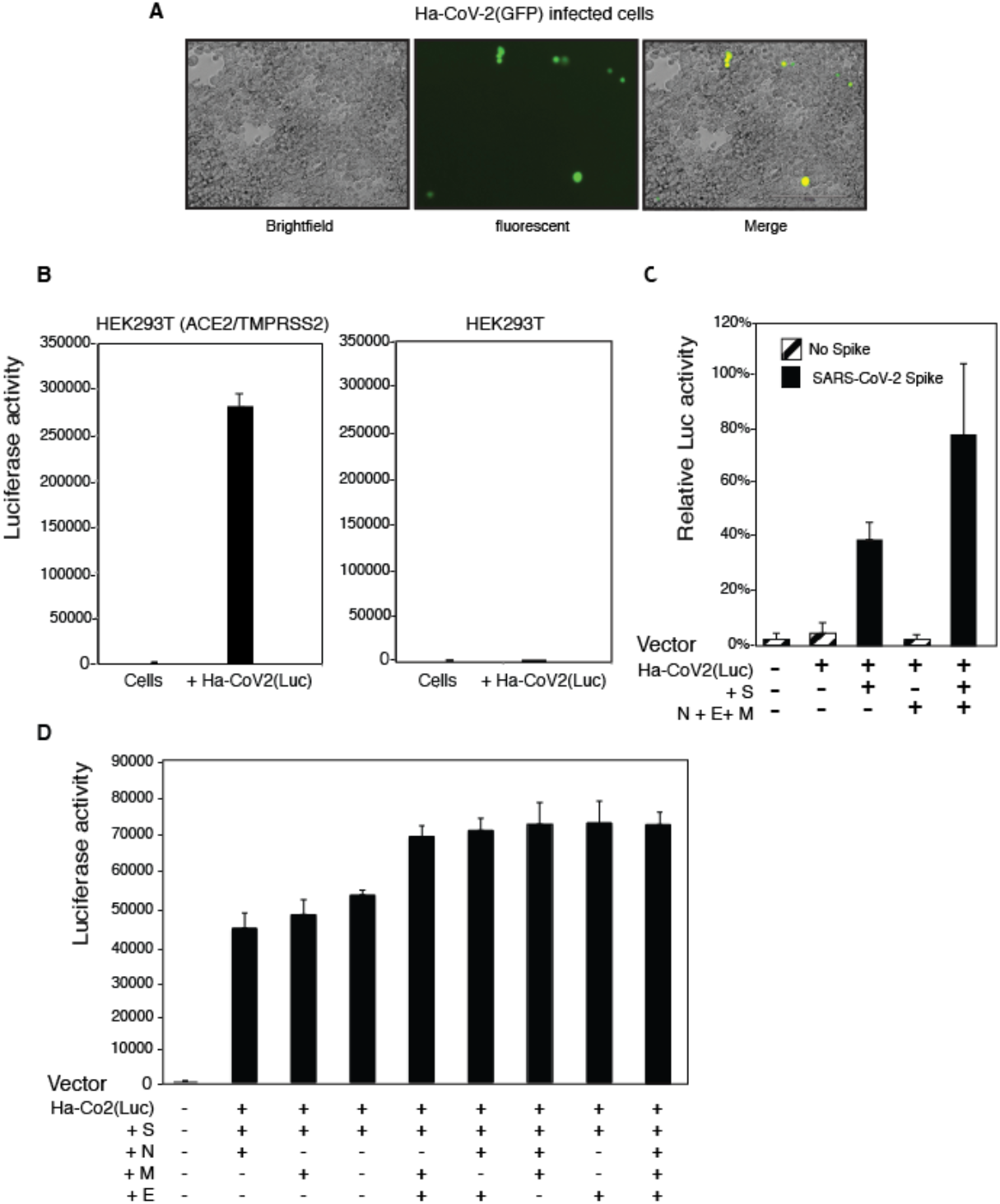
SARS-CoV-2 S protein and ACE2-dependent infection of target cells by Ha-CoV-2. **(A)** HEK293T(ACE2/TMPRSS2) cells were infected with Ha-CoV-2(GFP) particles. GFP expression was observed 48 hours post infection. **(B)** ACE2-dependent infection of target cells by Ha-CoV-2(Luc). HEK293T(ACE2/TMPRSS2) and HEK293T cells were infected with Ha-CoV-2(Luc) particles. Luciferase expression was quantified at 24 hours post infection. **(C)** SARS-CoV-2 S protein-dependent infection of target cells by Ha-CoV-2(Luc). Particles were assembled in the presence or absence of S or M + E + N. Luciferase expression was quantified at 4 hours post infection. (**D**) Requirements of M, E, and N for optimal infectivity of Ha-CoV(Luc). Particles were assembled in the presence S and combinations of individual proteins of M, E and N. Luciferase expression was quantified. Assays in **(B)** to **(D)** were performed in triplicates.

A major advantage of utilizing alphavirus-based RNA genome for Ha-CoV-2 is the extremely fast speed and high-level gene expression of alphaviruses; gene expression from the subgenomic RNA promoters occur within hours of infection, and levels of viral plus-RNAs can reach 200,000 copies in a single cell ^16,28^. We followed the time course of Ha-CoV-2(Luc) infection, and observed that the Luc reporter expression increased rapidly within 6 hours from the addition of particles to cells (**Fig. 3**). This rapid reporter expression kinetics permitted us to utilize Ha-CoV-2 for fast screening and quantification of neutralization antibodies and anti-viral drugs.

**Fig. 3.**
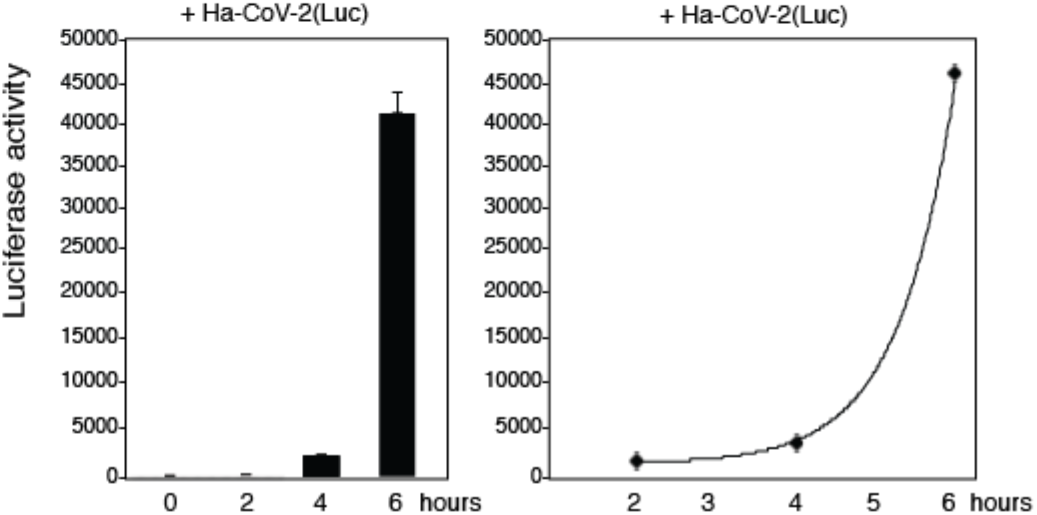
Rapid time course of reporter gene expression in Ha-CoV-2(Luc) infection. A 6 hour time-course of luciferase expression following infection of HEK293T(ACE2/TMPRSS2) cells with Ha-CoV-2(Luc) particles. Cells were infected with Ha-CoV-2(Luc) for 2 hours, washed, cultured in fresh medium, and then lysed and analyzed for Luc expression at different time points. The addition of virus to cells was defined as time “0”. Infection assays were performed in triplicates.

Lenti-based SARS-CoV-2 pseudoviruses have been commonly used for antiviral drug screening and neutralization antibody assays ^10,14^. We performed a comparison of Ha-CoV-2 with lenti-pseudovirus for the infection of both ACE2-overexpressing cells and cells expressing native levels of ACE2. Lenti-pseudovirus and Ha-CoV-2 particles were assembled in similar cell culture conditions, and an equal volume of the particles was used for infection. Both lenti-pseudovirus and Ha-CoV-2 can infect the ACE2-overexpressing HEK293T(ACE2/TMPRESS2) cells (**Fig. 4A**). However, infection of Calu-3, a human lung cancer cells expressing native levels of ACE2, was minimal with the lenti-pseudovirus ^31^, whereas Ha-CoV-2 particles produced much higher signal for the infection of Calu-3 cells (**Fig. 4B**). Infection of primary human ACE2-null monkey kidney cells with Ha-CoV-2 did not generate signals above uninfected cell background (**Fig. 4C**). These results demonstrate that Ha-CoV-2 is likely more sensitive for the infection of low ACE2-expressing cells.

**Fig. 4.**
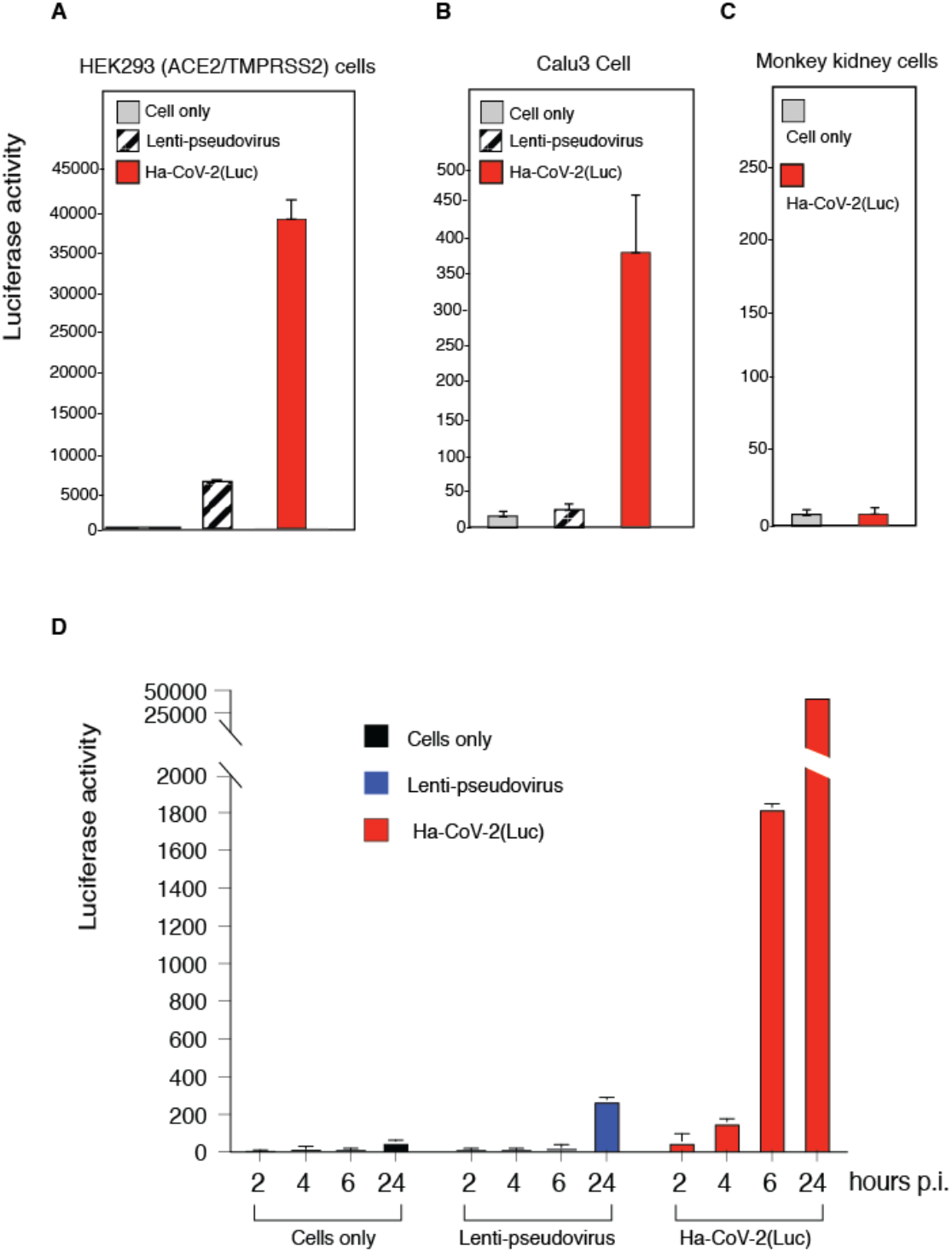
Comparison of SARS-CoV-2 S pseudotyped lentivirus with Ha-CoV-2 particles. **(A to C)**HEK293T(ACE2/TMPRSS2) and Calu-3 cells were infected with an equal volume of viral particles, Lenti-CoV-2(Luc) or Ha-CoV-2(Luc). Relative infection was quantified by luciferase assay at 72 hours post infection. Primary monkey kidney cells were also infected with Ha-CoV-2 for comparison. (**D**) Comparison of lenti-pseudovirus and Ha-CoV-2 in an infection time course. HEK293T(ACE2/TMPRSS2) were infected with an equal volume of viral particles, lenti-CoV-2(Luc) or Ha-CoV-2(Luc). Relative Luc reporter expression was quantified by luciferase assay from 2 to 24 hours post infection. All infection assays were performed in triplicates.

We further followed an infection time course of Ha-CoV-2, and compared it with the infection course of the lenti-pseudovirus. As shown in **Fig. 4D**, in Ha-CoV-2 infection, Luc reporter expression became detectable as early as 2 to 4 hours, whereas in the lenti-pseudovirus infection, Luc reporter expression was detectable only after 24 hours. In addition, the reporter expression in Ha-CoV-2 infection was much robust; by 24 hours, it reached a level approximately150 fold higher than that generated from the lenti-pseudovirus infection.

To validate Ha-CoV-2 for rapid screening and quantification of neutralizing antibodies, we tested an anti-SARS-CoV-2 antiserum (1F), which was serially diluted and pre-incubated with Ha-CoV-2(Luc). The antibody-virus complex was used to infect cells for 5 hours for Luc expression. As shown in **Fig. 5A**, we observed 1F concentration-dependent inhibition of Ha-CoV-2(Luc), and the IC50 was determined to be at 1:433 dilution (**Fig. 5A**). Given that SARS-CoV-2 lenti-pseudoviruses have been widely used in neutralization assays ^4,8,14^, we also performed a similar assay using 1F and a lenti-pseudovirus, Lenti-SARS-CoV-2(Luc) ^15^. Infected cells were analyzed at 72 hours post infection. We observed similar 1F concentration-dependent inhibition of the lenti-pseudovirus, and the IC50 was found to be at 1:186 dilution (**Fig. 5B**). These results demonstrated that Ha-CoV-2 is as effective as lenti-pseudoviruses for quantifying neutralizing antibodies, but with a much faster speed (5-12 hours versus 48-72 hours).

**Fig. 5.**
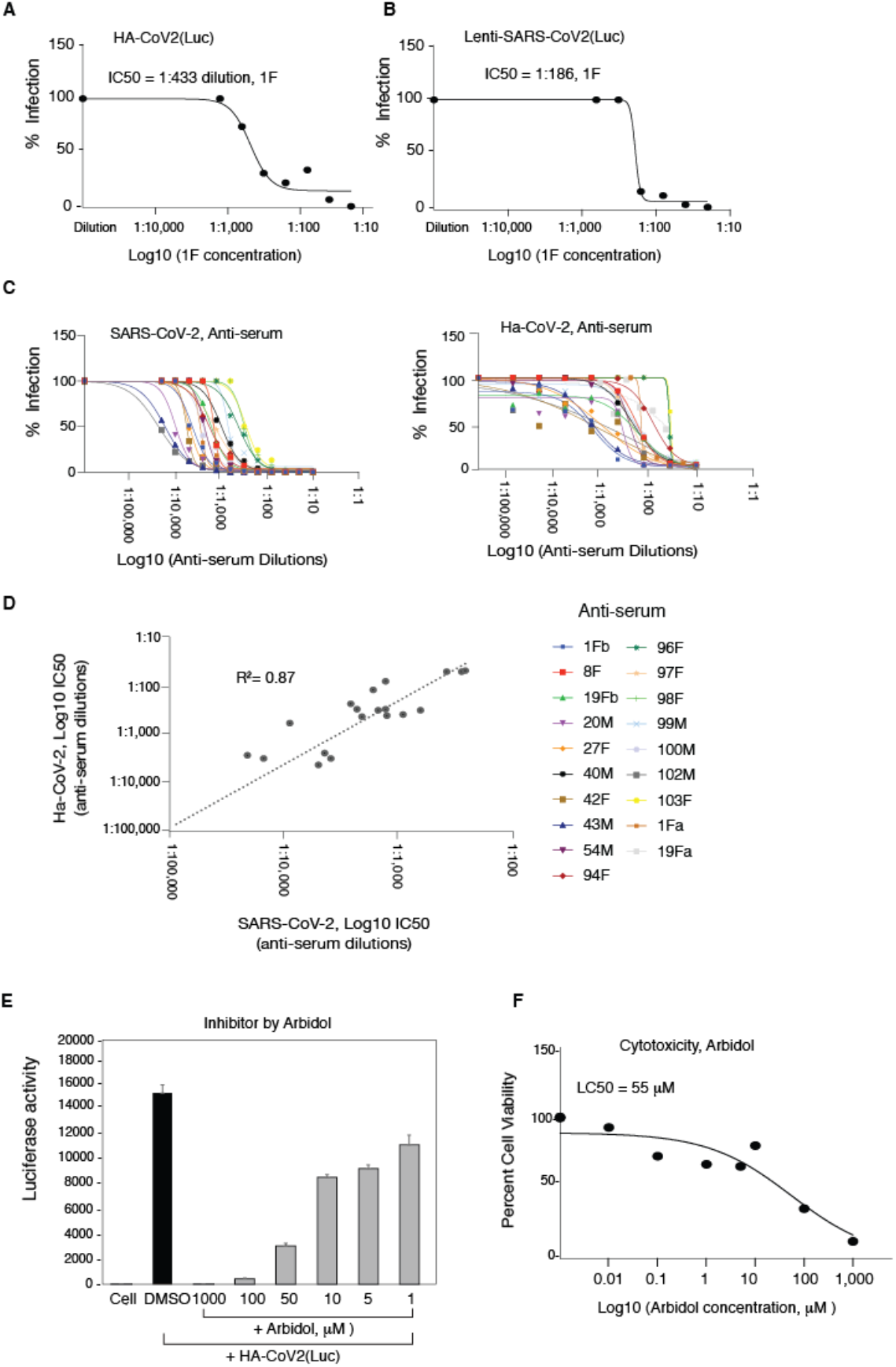
Validation of Ha-CoV-2 particles for rapid screening and quantification of neutralizing antibodies. (**A**) Quantification of neutralizing antibodies with Ha-CoV-2 particles. Shown are the concentration-dependent inhibition of Ha-CoV-2(Luc) by the anti-serum 1F and the 1F inhibition curve. 1F was serially diluted and incubated with Ha-CoV-2(Luc) particles for 1 hour at 37°C. The Ha-CoV-2(Luc)-antibody complex was used to infect HEK293T(ACE2/TMPRSS2) cells. Neutralization activities were quantified by luciferase assay at 5 hours post addition of virus to cells. Control serum was from healthy, uninfected donors. The IC_50_ was calculated using the relative percentage of infection versus serum concentration. For comparison, the anti-serum 1F was also similarly quantified using a SARS-CoV-2 S protein pseudotyped lentivirus, Lenti-CoV-2(Luc). Neutralization activities were quantified with luciferase assay at 72 hours post infection. (**C**and **D)**Correlation of serum neutralization activities quantified with Ha-CoV-2(Luc) and SARS-CoV-2. Convalescent plasma from 19 donors was quantified using infectious SARS-CoV-2 and plaque assays, or Ha-CoV-2(Luc). Neutralization activities were plotted and the IC_50_ values were calculated. The correlation in IC_50_ was plotted. (**E**and **F**) Rapid quantification of the anti-SARS-CoV-2 activity of Arbidol. HEK293T(ACE2/TMPRSS2) cells were pretreated for 1 hour with Arbidol. Cells were infected with Ha-CoV-2(Luc) in the presence of Arbidol. Viral entry inhibition was quantified by luciferase assay at 5 hours. An MTT cytotoxicity assay of Abidol was also performed on cells (**F**).

Based on the 1F results described above, we performed additional validation of Ha-CoV-2-based neutralizing assays using convalescent plasma from 19 donors. The inhibition curve and IC_50_ of each serum are presented in **Fig. 5C**. For comparison, an independent quantification was conducted using infectious SARS-CoV-2 to validate these anti-sera. We observed a direct correlation (r^2^ = 0.87) in the IC_50_ values obtained from Ha-CoV-2 and from SARS-CoV-2 (**Fig. 5D**). These results demonstrated that Ha-CoV-2 can be used for rapid quantification of neutralizing antibodies.

Pseudoviruses have also been commonly used for high throughput screening of SARS-CoV-2 entry inhibitors ^7,10^. We tested a broad-spectrum viral entry inhibitor, Arbidol (Umifenovir) ^32^, for its ability to block Ha-CoV-2(Luc) infection. As shown in **Fig. 5E**, we observed dosage-dependent inhibition of Ha-CoV-2(Luc) in 5 hours, and the IC_50_ was determined to be 16 μM. These results demonstrated that Ha-CoV-2 can be used for rapid screening of SARS-CoV-2 entry inhibitors.

Finally, we investigated whether the Ha-CoV-2 system can be used for rapid evaluation of relative infectivity of viral variants. The D614G spike mutation emerged early in the COVID-19 pandemic, and has recently been reported to confer greater infectivity that has led to the global dominance of the D614G mutant in circulation ^33,34^. To determine whether the increase in virus infectivity can be recapitulated and quantified by the Ha-CoV-2 system, we assembled Ha-CoV-2 particles using the G614 mutant S protein (G614) or the parental S protein (D614). We found that the D614G mutation did not increase virion release or the level of S protein virion incorporation (**Fig. 6A**and **6B**). However, Ha-CoV-2 particles bearing the G614 spike were found to be nearly 3 times more infectious than those bearing the D614 spike (**Fig. 6C**).

**Fig. 6.**
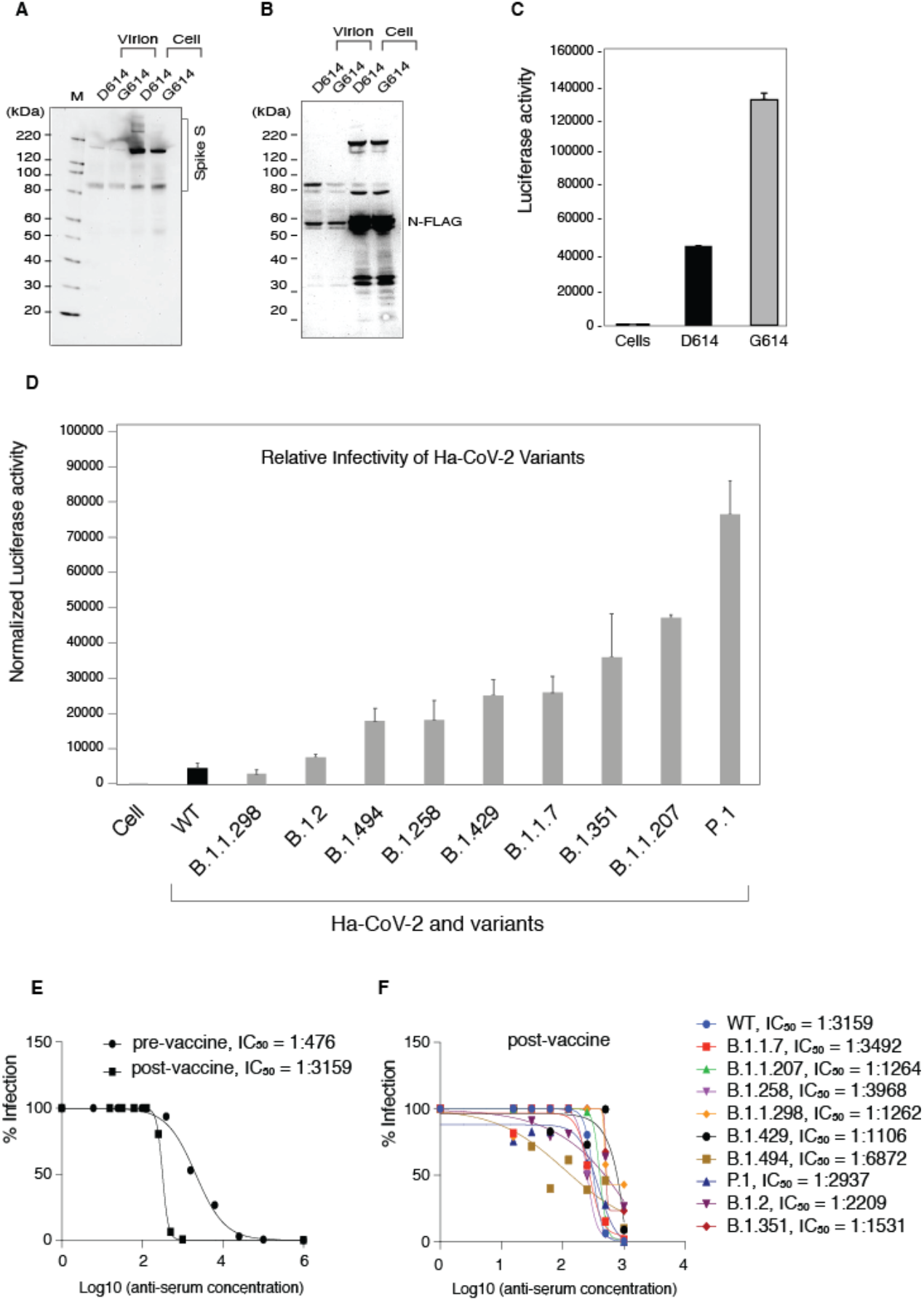
Quantification of the relative infectivity of Ha-CoV-2 variants and their responses to neutralizing antibodies. (**A** and **B**) Ha-CoV-2(Luc) particles bearing the G614 mutation S or the parent D614 S were assembled, and analyzed for the incorporation of S and N in virions. (**C**) Ha-CoV-2(Luc)(G614) or Ha-CoV-2(D614) was used to infect target cells, and Luc expression was quantified at 5 hours. An equal level of viral particles was used for infection. Infection assays were performed in triplicates. (**D**) A panel of 9 S protein mutants from SARS-CoV-2 variants were used to assemble Ha-CoV-2(Luc) particles, and then used to infect target cells. The relative infectivity was quantified and normalized with the genomic RNA copies of individual Ha-CoV-2(Luc) variants. WT refers to Ha-CoV-2 derived from the original SARS-CoV-2 strain. Infection assays were performed in triplicates. (**E**and **F**) Quantification of anti-serum against Ha-CoV-2(Luc) and its variants. Convalescent plasma from an infected blood donor, before and after one dose vaccination, was quantified for inhibition of Ha-CoV-2(Luc) infection. Neutralization activities were quantified by luciferase assay at 12 hours post infection. The IC_50_ was calculated using the relative percentage of infection versus serum concentration (**E**). The post-vaccination anti-serum was similarly quantified for the inhibition of Ha-CoV-2(Luc) variants (**F**).

We further assembled additional 9 Ha-CoV-2(Luc) isolates derived from circulating SARS-CoV-2 variants (selected from the GISAID global reference database, **Table 1**), including the Brazil variant (P.1), the South Africa variant (B.1.351), the UK variant (B.1.1.7), the California variant (B.1.429), and several other emerging variants (B.1.2, B.1.494, B.1.1.207, B.1.258, and B.1.1.298). Ha-CoV-2(Luc) and the related S protein variants were used to infect target cells, and the relative infectivity was quantified. As shown in **Fig. 6D**, when normalized with the genomic RNA copies, these variants in general are 2-10 fold more infectious than the original parental Ha-CoV-2(Luc). These results demonstrated that Ha-CoV-2 can provide a convenient tool for rapid monitoring and quantification of viral variants. As a proof-of-concept, we further quantified the ability of an anti-serum to neutralize viral variants. We acquired convalescent plasma from a donor who was infected, and then vaccinated with one dose Moderna mRNA-1273. This one dose vaccination has greatly increased the anti-serum titer for approximately 6 fold (**Fig. 6E**). Furthermore, when Ha-CoV-2(Luc) variants were tested, we found that the post-vaccination serum had neutralizing activities against all variants tested (**Fig. 6F**). Nevertheless, the neutralizing activities differ greatly among the variants; the anti-serum has the highest neutralizing activity against B.1.494 (IC_50_, 1: 6872), and lowest activity against the B.1.1.429 variant (IC_50_, 1:1106). These results demonstrate that Ha-CoV-2 can be used for rapid quantification of SARS-CoV-2 variants for potential impacts on neutralizing antibodies and vaccine effectiveness

**Table 1.** List of S protein mutations in SARS-CoV-2 isolates.

| **GISAID**<br>**Variant lineage** | **Mutations in S protein** | **GISAID**<br>**Accession number** |
| --- | --- | --- |
| B.1.1.7 | A570D, D614G, D1118H, H69del, N501Y, P681H, S982A, T716I, V70del, Y145del | EPI_ISL_581117 |
| B.1.1.207 | D614G, E484K, P681H | EPI_ISL_778908 |
| B.1.1.298 | D614G, H69del, I692V, M1229I, V70del, Y453F | EPI_ISL_616802 |
| B.1.2 | D614G, E484K, G446V, Y453F | EPI_ISL_833413 |
| B.1.258 | D614G, H69del, N439K, V70del | EPI_ISL_755592 |
| B.1.351 | A243del, A701V, D80A, D215G, D614G, E484K, K417N, L242del, L244del, N501Y | EPI_ISL_678597 |
| B.1.429 | A222V, D614G, L452R, S13I, W152C | EPI_ISL_847764 |
| B.1.494 | A262S, D614G, D796Y, H49Y, L452R, N501Y, P681R, Q613H | EPI_ISL_826591 |
| P.1 | D138Y, D614G, E484K, H655Y, K417T, L18F, N501Y, P26S, R190S, T20N, T1027I, V1176F | EPI_ISL_833136 |

\* SARS-CoV-2 lineage identification and variant naming were obtained from GISAID (https://www.gisaid.org); mutations in the spike protein and the sequence accession number of each isolate are listed.

## DISCUSSION

The study of SARS-CoV-2 requires high-level containment that limits the use of infectious virus in common clinical and research laboratories. Pseudoviruses and virus-like particles (VLPs) have been widely used for SARS-CoV-2 drug discovery and vaccine development. Pseudoviruses, such as those derived from lentivirus and vesicular stomatitis virus, can mimic the entry process of SARS-CoV-2. However, structurally, they are very different from SARA-CoV-2 and lack structural components provided by M, E, and N of SARS-CoV-2. VLPs closely resemble SARS-CoV-2 particles, but VLPs contain no genome for reporter expression in target cells ^35^. In this article, we described the development and validation of a novel hybrid system, the Ha-CoV-2 particle, which is structurally a VLP, but possesses the ability of a pseudovirus to enter and express reporter genes in target cells. The genome of Ha-CoV-2 is derived from alphavirus, which allows for rapid and robust quantification of reporter expression within hours of viral entry. We further demonstrated that Ha-CoV-2 can be used for rapid screening and quantification of neutralization antibodies, viral variants, and antiviral drugs.

We also performed a direct comparison between Ha-CoV-2 and a lenti-pseudovirus in antibody neutralization assays. While both systems are effective in quantifying neutralizing antibodies, the sensitivity of the two systems differ. The lenti-pseudovirus contains only the S protein of SARS-CoV-2, whereas Ha-CoV-2 contains all four structural proteins (S, M, E, and N) of SARS-CoV-2, and has no structural proteins from other viruses (e.g. *gag* and *pol* of lentivirus). Although S is the primary requirement for viral entry, the presence of other structural proteins of SARS-CoV-2 may also affect virion infectivity and particle interaction with cell membrane and antibodies. In our system, the lack of M and E on virion particles does appear to affect virus infection (**Fig. 2D**).

In addition to viral structural proteins, virion particles also incorporate multiple cellular proteins during virion budding and release. Many of these cellular factors such as PSGL-1 can impact virion infectivity ^15,36–38^ and antibody binding to plasma membrane ^39^. SARS-CoV-2 budding occurs mainly at the membranes of ER-Golgi intermediate compartment ^40^, whereas the lenti-pseudovirus buds from the plasma membrane ^41^. Because of the difference, it is possible that different sets of cellular proteins may be incorporated into lenti-pseudovirus and SARS-CoV-2. In this regard, the close resemblance of Ha-CoV-2 particle to SARS-CoV-2 may provide a unique tool for studying effects of virion host proteins in SARS-CoV-2 infection and pathogenesis ^15^.

As SARS-CoV-2 continues to circulate and evolve, viral variants pose a particular challenge for the control of the COVID-19 pandemic, as documented in the recent emergence of the B.1.1.7 lineage in UK ^42^. Viral mutation may lead to increases in viral transmission and fitness, and thus there is an urgent need for rapid identification and characterization of emerging variants for changes in viral infectivity and responses to neutralizing antibodies. The Ha-CoV-2 system would provide a robust platform for rapid quantification of viral variants and potential impacts on neutralizing antibodies and vaccine effectiveness.

## Materials and Methods

### Virus and viral particle assembly

The SARS-CoV-2 S, S(D614G), M, E, or N expression vectors were purchased from Sinobiological. The Ha-CoV-2(Luc) and Ha-CoV-2(GFP) vectors, and the S protein variants were selected from isolates identified in the GISAID global database (Table 1), and synthesized by Twist Bioscience. Ha-CoV-2 particles were assembled by cotransfection of HEK293T cells in 10 cm dish with 2.5 μg of each of the SARS-CoV-2 structural protein expression vectors (S, N, E, M) and 10 μg of Ha-CoV-2(Luc) or Ha-CoV-2(GFP). Particles were harvested at 48 hours post cotransfection, filtered through a 0.45 μm filter, and then concentrated by gradient centrifugation. Lenti-pseudovirus was assembled by cotransfection of HEK293T cells with SARS-CoV-2 S expression vector (0.5 μg), pCMV∆R8.2 (7.5 μg), and pLTR-Tat-IRES-Luc (10 μg) as previously described ^15^.

### Detection of Ha-CoV-2 virion incorporation of structural proteins

The SARS-CoV-2 M-FLAG and N-FLAG vectors were kindly provided by Dr. Pei-Hui Wang ^43^. HEK293T cells were co-transfected with 10 μg Ha-CoV-2(Luc), 2.5 μg of the SARS-CoV-2 S expression vector, and 2.5 μg each of the M-FLAG, N-FLAG, and E-FLAG vectors. Particles were harvested, filtered through a 0.45 μm filter, and then purified by gradient centrifugation. Virion lysates were analyzed by SDS-PAGE and western blot using Spike Protein S2 Monoclonal Antibody (1A9) (Invitrogen) (1:1000 dilution) or DYKDDDDK Tag Monoclonal Antibody (FG4R) (Invitrogen) (1: 1000 dilution). Membranes were then incubated with Anti-mouse IgG, HRP-linked Antibody (Cell signaling) (1: 2000 dilution) for 60 min at room temperature. Chemiluminescence signal was detected by using West Pico or West Femto chemiluminescence reagent (Thermo Fisher Scientific). Images were captured with a CCD camera (FluorChem 9900 Imaging Systems) (Alpha Innotech). Particles were also captured with magnetic beads for analyses. Briefly, magnetic Dynabeads Pan Mouse IgG (Invitrogen) (2×10^7^ beads/50 μl) were conjugated with Spike Protein S2 Monoclonal Antibody (1A9) (Invitrogen) (2 μl antibody) for 30 minutes at room temperature. After conjugation, virions were incubated with the anti-S2-beads for 30 minutes at 4°C, and pulled down with a magnet. After washing with cold PBS for 5 times, virions were lysed in LDS lysis buffer (Invitrogen). Lysates were analyzed by SDS-PAGE and western blot using DYKDDDDK Tag Monoclonal Antibody (FG4R) (Invitrogen) (1: 1000 dilution) to detect FLAG-Tagged SARS-CoV-2 M proteins.

### Viral infectivity assay

Ha-CoV-2 particles were used to infect HEK293T(ACE2/TMPRSS2) cells (a gift from Virongy LLC, Manassas, VA), Calu-3 cells (ATCC), HEK293T cells (ATCC) and primary monkey kidney cells provided by Dr. Xuefeng Liu. Briefly, cells were seeded in 12-well plates (2×10^5^ cells) per well. Cells were infected for 1-2 hours at 37°C, washed, cultured in fresh medium for 3-48 hours, and then lysed in Luciferase Assay Lysis Buffer (Promega) for luciferase activity using GloMax Discover Microplate Reader (Promega). Lenti-pseudovirus particles were used to infect HEK293T(ACE2/TMPRSS2) cells and Calu-3 cells (ATCC). Cells were infected for 2 hours, cultured for 3 days, and then lysed in Luciferase Assay Lysis Buffer (Promega) for luciferase assays using GloMax Discover Microplate Reader (Promega).

### Neutralizing Antibody Assay

Ha-CoV-2 particles were pre-incubated with serially diluted sera from COVID19 patients for 1 hour, and then added to HEK293T(ACE2/TMPRSS2) cells for 2 hours. Cells were then washed, and cultured in fresh medium for additional 3-24 hours. Cells were lysed in Luciferase Assay Lysis Buffer (Promega) for luciferase assays using GloMax Discover Microplate Reader (Promega). For neutralization assays using wild-type SARS-CoV-2 virus, anti-serum was serially diluted (a twelve-point, two-fold dilution series starting at 1:10 dilution), and pre-incubated with 100 pfu of SARS-CoV-2 for 1 hour at 37 °C. After incubation, viral plaque assay was conducted to quantify viral titers. Briefly, Vero cells (ATCC) in 12-well plates (2×10^5^ cells per well) were infected with virus for 1 hour at 37 °C. After infection, a 1:1 overlay, consisting of 0.6% agarose and 2X EMEM without phenol red (Quality Biological), supplemented with 10% fetal bovine serum (FBS) (Gibco), was added to each well. Plates were incubated at 37°C for 48 hours. Cells were fixed with 10% formaldehyde for 1 hour at room temperature, and then the agarose overlay was removed. Cells were stained with crystal violet (1% CV w/v in a 20% ethanol solution). Viral titer of SARS-CoV-2 was determined by counting the number of plaques.

### Antiviral Drug Assay

Arbidol-hydrochloride (Sigma) was resuspended in Dimethyl sulfoxide (Sigma). HEK293T(ACE2/TMPRSS2) cells were pretreated for 1 hour with serially diluted Arbidol. Ha-CoV-2 particles were added cells, followed by the addition of Abidol to maintain the drug concentration. Cells were infected in the presence of Arbidol for 2 hours, washed, and then cultured in fresh medium for a total of 5 hours. Cells were lysed in Luciferase Assay Lysis Buffer (Promega) for luciferase assays using GloMax Discover Microplate Reader (Promega).

## ACKNOWLEDGMENTS

The authors wish to thank Janice Yoon for technical assistance. Dr. Pei-Hui Wang for providing FLAG-tagged M, N, and E expression vectors. This work was supported by George Mason University internal research fund.

## AUTHOR CONTRIBUTIONS

Experiments were designed by B. H. and Y.W. Manuscript was written by Y.W. Experiments were performed by B.H., S.H., L.D.C., D.D., F.A., A.N., A.L., T.L., X.L., and L.L.

## Data availability

All data generated or analyzed during this study are included in this article.

